# The ‘phruta’ R package and ‘salphycon’ shiny app: increasing access, reproducibility, and transparency in phylogenetic analyses

**DOI:** 10.1101/2023.01.11.523621

**Authors:** Cristian Román-Palacios

## Abstract

1. Current practices for assembling phylogenetic trees often recur to sequence data stored in GenBank. However, the molecular and taxonomic make up of sequences deposited in GenBank is generally not very clear.
2. ‘phruta’, a newly developed ‘R’ package, is designed to primarily improve access to genetic data stored in GenBank. Functions in ‘phruta’ further enable users to assemble single- and multi-gene molecular datasets, and run basic phylogenetic tasks, all within ‘R’.
3. The structure of the functions implemented in ‘phruta’, designed as a workflow, aim to allow users to assemble simple workflows for particular tasks, which are in turn expected to increase reproducibility of relatively simple phylogenies.
4. To support the use of ‘phruta’ by researchers in different fields with variable levels of coding expertise, this paper presents ‘salphycon’, a shiny web app that is expected to increase access to the fundamental functions in the ‘phruta’ ‘R’ package.

## 1 INTRODUCTION

Although the assemblage of molecular phylogenies has been the backbone of numerous studies, the existing tools used to retrieve sequences from public databases, curate molecular datasets, assemble multi-locus alignments, and finally infer phylogenies generally involves an extensive set of software that, in many cases, is poorly streamlined (Hall, 2004; Lemey et al., 2009; Wiley, and Lieberman, 2011). Here, I present ‘phruta’, an ‘R’ package, and ‘salphycon’, the associated interactive application, designed to improve the reproducibility of and access to a fraction of the existing tools of the phylogenetic workflow.

While similar functions for assembling curated molecular datasets for phylogenetic analyses can be found in ‘phylotaR’ (Wang et al. 2022) and ‘SuperCRUNCH’ (Portik and Wiens 2020), ‘phruta’ simplifies into a single open-source package the numerous required tools. For example, ‘phylotaR’ is limited to downloading and curating sequences (e.g. it does not align sequences) and, ‘SuperCRUNCH’ curates sequences that are already stored locally. ‘SUPERSMART’ (Scharn et al. 2016; Antonelli et al. 2017) and the associated ‘R’ workflow ‘SUPERSMARTR’ (Antonelli et al. 2017; Bennett et al. 2018) also contain similar functions, some of which have been simplified here in a single package, ‘phruta’. ‘phruta’ improves upon standalone applications ‘MEGA’ (Tamura et al. 2007) and geneious (Kearse et al. 2012) by decreasing the burden for researchers to perform and publish reproducible computational workflows (e.g. ‘MEGA’) and eliminating unnecessary financial barriers for all researchers in performing said analyses (e.g. geneious has a paid version). Finally, note that ‘phruta’ is already integrated within ‘R’ (R Core Team, 2022), a language that contains the state of art packages and methodological implementations in phylogenetics and many different fields in biology and related fields (e.g. Revell and Harmon, 2022).

In short, ‘phruta’ and ‘salphycon’ provide an early perspective on how to improve the reproducibility and access to the phylogenetic pipeline in ‘R’. In this manuscript, I provide a brief overview of ‘phruta’, briefly outlining its current capabilities. Finally, I present ‘salphycon’, a Shiny app (Chang et al. 2021) designed to even further increase access to reproducible phylogenetic pipelines via’phruta’, and discuss its functionality and future outlook.

## 2 THE ‘phruta’ ‘R’ PACKAGE

The ‘phruta’ package is primarily designed to simplify the basic phylogenetic computational workflow, entirely in ‘R’. ‘phruta’ is expected to allow scientists to assemble molecular datasets or phylogenies for particular taxonomic groups, with minimal complexity and maximal reproducibility.

In general, ‘phruta’ (1) scans GenBank (Benson et al. 2012) primarily using the ‘rutils’ (Schöfl 2016) and ‘rentrez’ (Winter, 2017) ‘R’ packages to identify potential phylogenetically relevant gene regions for a given set of taxa, (2) retrieves gene sequences and curates taxonomic information from different taxonomic backbones, (3) combines downloaded and local gene sequences, and (4) performs phylogenetic tasks such as sequence alignment, phylogenetic inference, and basic tree dating. An interactive web application, ‘salphycon’, is also available as an additional resource to run the basic functions of ‘phruta’ without the need for any previous ‘R’ programming experience.

The current release of ‘phruta’ includes a set of eight major functions. Depending on the information available, running all eight major functions in ‘phruta’ potentially results in a time-calibrated phylogeny for a given set of taxa. Note that all the functions for which the primary output is sequences (aligned or unaligned) are listed under ‘sq.*’. All the functions that output phylogenies (time-calibrated or not) are listed under ‘tree.*’.

- First, the distribution of genes sampled in GenBank for a given organism or set of taxa can be explored using the ‘acc.gene.sampling()’ function. This function in ‘phruta’ extends the functionality of the ‘rutils’ ‘R’ package by scraping data from GenBank. ‘acc.gene.sampling()’ returns a table summarizing the distribution of genes sampled for a given taxon or set of taxa. Note that ‘acc.gene.sampling()’ only retrieves a set of gene names but does not download any sequences.
- Second, a list of accession numbers for given a list of organisms can be retrieved using the ‘acc.table.retrieve()’ function. Instead of directly downloading sequences from GenBank (see ‘sq.retrieve.direct()’ below), retrieving accession numbers as an intermediate step, allows users to have more control over the sequences that are being used in downstream analyses.
- Third, two different functions in ‘phruta’ enable users to download sequences from GenBank. First, sequences can be downloaded using ‘sq.retrieve.indirect()’ on the accession numbers retrieved before using the ‘acc.table.retrieve()’ function presented above. Second, users can also skip defining an accession number table and download gene sequences directly using the ‘sq.retrieve.direct()’ function in ‘phruta’. Note that ‘sq.retrieve.direct()’ is primarily based on functions from ‘rentrez’ (Winter, 2017) and ‘sq.retrieve.indirect()’ mainly uses functions from ‘rutils’ (Schöfl 2016). Optionally, these functions will create a folder ‘0.Sequences’ that includes all the retrieved sequences. Note that using ‘acc.table.retrieve()’ is the preferred option within ‘phruta’.
- Fourth, local sequences to those retrieved from GenBank can be included into the workflow using the ‘sq.add()’. This function saves all resulting ‘fasta’ files in two directories: (1) combined sequences are stored in ‘0.Sequences’ and local sequences are moved into ‘0.AdditionalSequences’. Note that the originally downloaded sequences are moved to ‘0.0.OriginalDownloaded’ at this step. ‘sq.add()’ is primarily based on functions in the ‘ape’ ‘R’ package (Paradis and Schliep, 2019).
- Fifth, the ‘sq.curate()’ function filters out unreliable sequences based on information listed in GenBank (e.g. the prefix ‘PREDICTED’) and on the target taxonomic information provided by the user. For instance, if a given species belongs to a non-target group, this species is dropped from the analyses. ‘sq.curate()’ function automatically corrects taxonomy based on alternative taxonomic databases, renames sequences, and further uses the ‘odseq’ ‘R’ package (Jiménez, 2022) to detect outliers in the set of sequences.
- Sixth, ‘sq.aln()’ performs multiple sequence alignment in ‘fasta’ files. ‘phruta’ uses the ‘DECIPHER’ ‘R’ package for this purpose (Wright 2016, 2020), which allows for adjusting sequence orientation and masking (removing ambiguous sites).
- Seventh, the ‘tree.raxml()’ function allows users to perform tree inference under ‘RAxML’ for sequences in a given folder. This is a wrapper to ‘ips::raxml()’ (Heibl, 2008) and each of the arguments can be customized. The current release of ‘phruta’ can manage both partitioned and unpartitioned analyses in RAxML (Stamatakis, 2014). Note that users can also provide starting and constrained trees.
- Eight, ‘tree.dating()’ enables users to perform time-calibrations of a given phylogeny (Eastman et al. 2013) using ‘geiger::congruify.phylo()’ (Harmon et al. 2008; Pennell et al., 2014). ‘phruta’ includes a basic set of comprehensively sampled, time-calibrated phylogenies that are used to extract secondary calibrations for the target phylogeny. Users can choose to run either ‘PATHd-8’ (Britton et al. 2007) or ‘treePL’ (Smith and O’Meara, 2012) for the calibration step.

## 3 BRIEF TUTORIAL: USING ‘phruta’ TO INFER THE PHYLOGENETICS OF NEW WORD QUAILS

Let’s learn how ‘phruta’ works by assembling a molecular dataset at the species level for a handful of bird genera. Note that although this tutorial is based on a particular set of taxa, users can decide on what can choose their target clades in other families, orders, or even kingdoms.

Here, we will focus on assembling a phylogeny for the new world quail (Johnsgard, 1998). Species in this group are classified in the family Odontophoridae, a clade including nearly 34 extant species classified in 10 genera. In general, the higher-level taxonomic information in GenBank for the Odontophoridae is largely congruent with recent studies on the systematics of the group (Crowe et al. 2006a, b; Cohen et al. 2012; Hosner et al. 2015). However, GenBank classifies *Ptilopachus*, a genus commonly included under the Odontophoridae, as part of the Phasianidae. We will follow more recent studies suggesting that *Ptilopachus* is nested within the Odontophoridae. As our outgroup, we will select the Phasianidae. Within this later family, Phasianidae, we will explicitly focus on sampling species in the genus *Polyplectron*, a clade of eight extant species. Finally, given that the systematics of the Odontophoridae has been discussed before using morphological and molecular evidence, we will be able to compare the topology of our tree relative to recent studies (Crowe et al. 2006a, b; Cohen et al. 2012; Hosner et al. 2015).

Up to this point, we have decided the taxonomic makeup of our analyses. From here, we could simply check the genetic sampling used in previous studies and search for those genes in GenBank for Odontophoridae and *Polyplectron* (Crowe et al. 2006a, b; Cohen et al. 2012; Hosner et al. 2015). Alternatively, we could use ‘phruta’ to determine genes well sampled in GenBank for both the ingroup and outgroup. For simplicity, we will follow the latter procedure by using the ‘gene.sampling.retrieve()’ function in ‘phruta’. An object of class ‘data.frame’ named ‘gs.seqs’ is generated containing the names of different gene regions that are sampled in GenBank for the target taxa.

~~~
‘“{r}
gs.seqs <- gene.sampling.retrieve(organism = c(“Odontophoridae”, “Ptilopachus”,
“Polyplectron”), speciesSampling = TRUE, npar = 6, nSearchesBatch = 500)
”’
~~~

Given the search terms, ‘phruta’ retrieved the names for ~80 gene regions from GenBank. Note that the ‘gene.sampling.retrieve()’ function provides an estimate of the number of species in GenBank that match the taxonomic criteria of the search term and that have sequences for a given gene region. However, the distribution of gene names and their specieslevel coverage is only as good as the annotations for genes deposited in GenBank. I show a summary of the resulting search in Table 1.

**Table 1.** Top six gene regions sampled in GenBank for the Odontophoridae and *Polyplectron*. In the table, the number of species sampled for a given gene region is shown along with the relative number of species in GenBank across the examined genes. Note that ‘phruta’ was able to retrieve the names of ~80 gene regions but only a summary of them are shown in the table.

| Gene | Sampled in # of<br>species | Percent of sampled<br>species |
| --- | --- | --- |
| NADH dehydrogenase subunit 2 | 92 | 99 |
| 12S ribosomal RNA | 28 | 30 |
| eukaryotic elongation factor 2 | 26 | 28 |
| NADH dehydrogenase subunit 5 | 19 | 20 |
| cytochrome b | 16 | 17 |
| cytochrome oxidase subunit 1 | 10 | 11 |

Using the combination of well-sampled genes and our list of taxa, we will now generate a preliminary summary of the accession numbers. I call this dataset preliminary because not all these accession numbers are expected to be in the final molecular dataset. For instance, some sequences may be removed after taxonomic synonyms are identified in the dataset.

From this point, we will assemble a species-level summary of accession numbers using the ‘acc.table.retrieve()’ function in ‘phruta’ (i.e. ‘speciesLevel = TRUE’ argument). For simplicity, this tutorial will focus on analyzing gene regions that are sampled in >20% of the species (‘targetGenes’ data.frame). The ‘acc.table’ object created below is a ‘data.frame’ object that will later be used to download the relevant gene sequences from GenBank (see Table 2).

~~~
‘“{r}
targetGenes <- gs.seqs[gs.seqs$PercentOfSampledSpecies > 20,]
acc.table <- acc.table.retrieve(
        clades = c(“Odontophoridae”, “Ptilopachus”, “Polyplectron”),
        genes = targetGenes$Gene,
        speciesLevel = TRUE,
        npar = 6,
        nSearchesBatch = 500
       )
”’
~~~

**Table 2.**
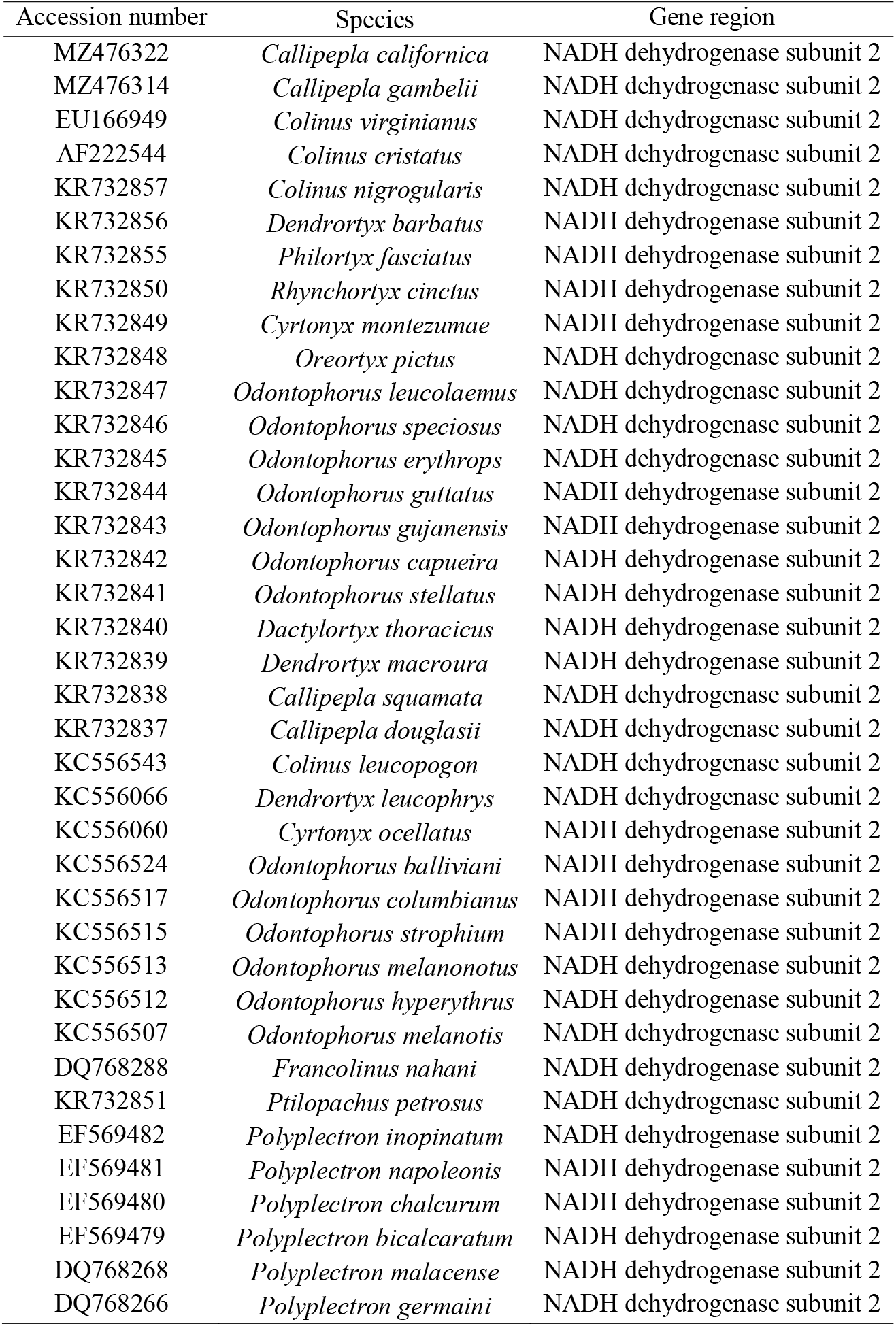
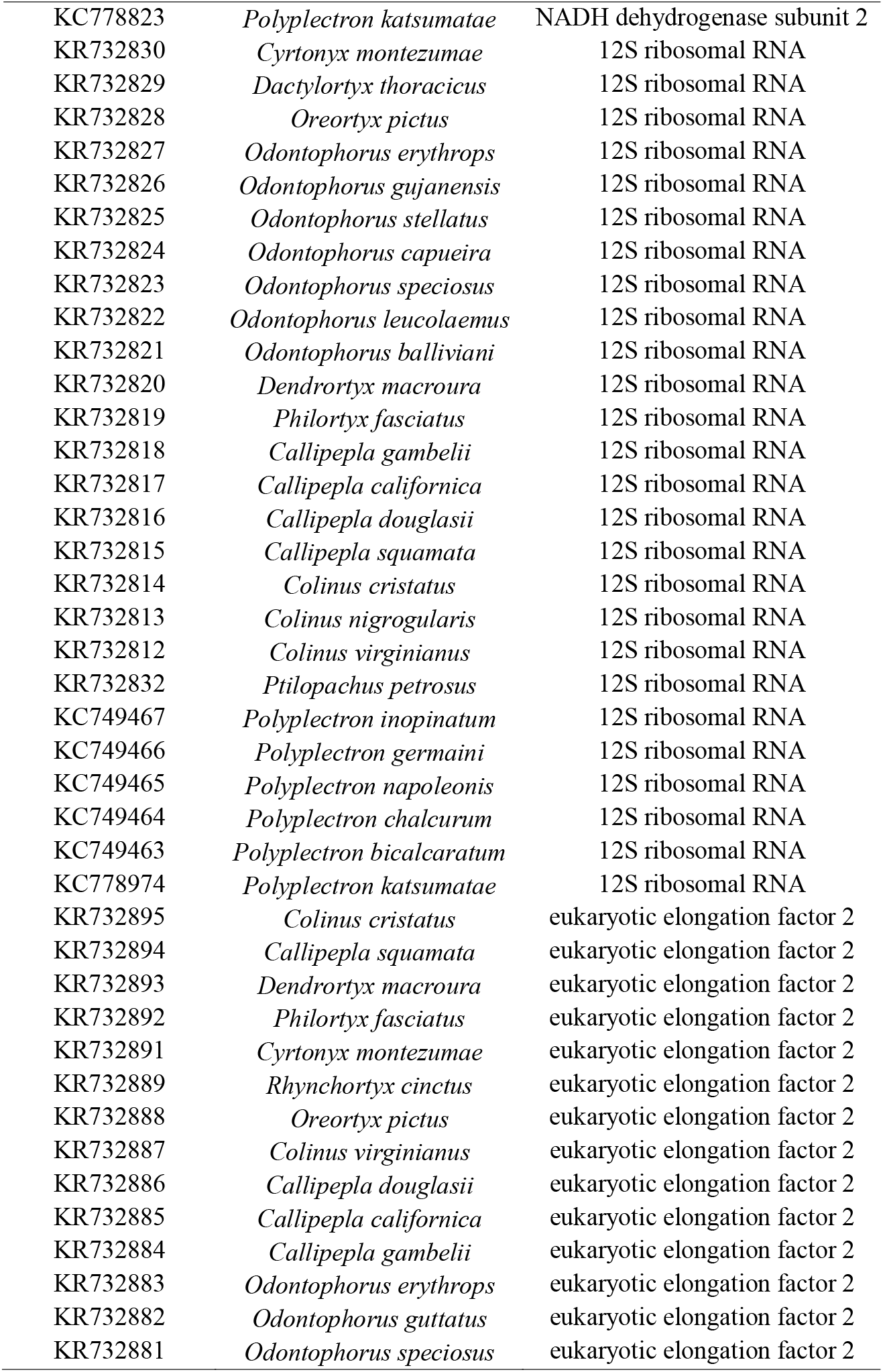
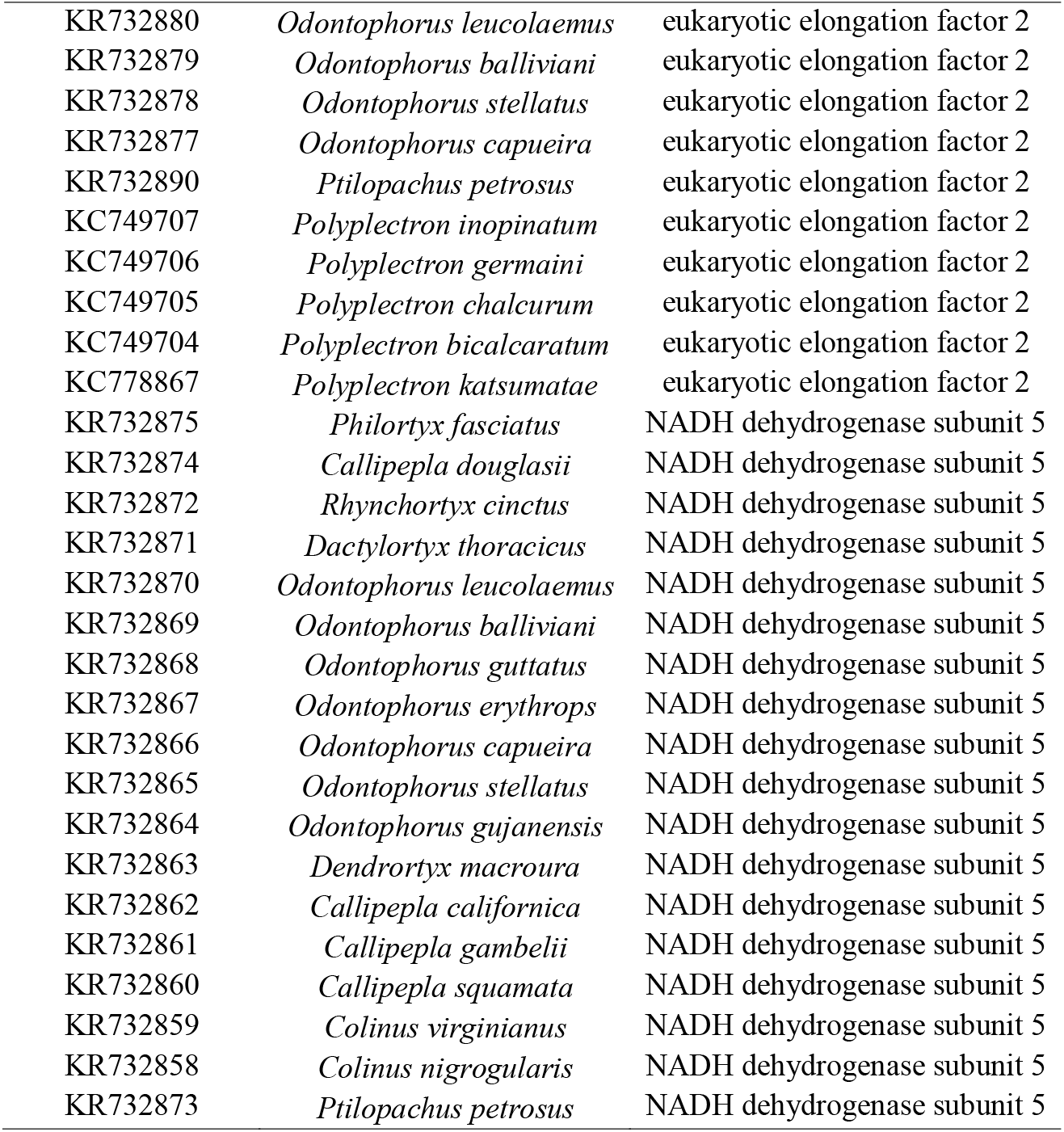
Accession numbers for the Odontophoridae and *Polyplectron* obtained using ‘phruta’. We show the species-level sampling based on gene regions with >20% of species (see Table 1).

| Accession number | Species | Gene region |
| --- | --- | --- |
| MZ476322 | *Callipepla californica* | NADH dehydrogenase subunit 2 |
| MZ476314 | *Callipepla gambelii* | NADH dehydrogenase subunit 2 |
| EU166949 | *Colinus virginianus* | NADH dehydrogenase subunit 2 |
| AF222544 | *Colinus cristatus* | NADH dehydrogenase subunit 2 |
| KR732857 | *Colinus nigrogularis* | NADH dehydrogenase subunit 2 |
| KR732856 | *Dendrortyx barbatus* | NADH dehydrogenase subunit 2 |
| KR732855 | *Philortyx fasciatus* | NADH dehydrogenase subunit 2 |
| KR732850 | *Rhynchortyx cinctus* | NADH dehydrogenase subunit 2 |
| KR732849 | *Cyrtonyx montezumae* | NADH dehydrogenase subunit 2 |
| KR732848 | *Oreortyx pictus* | NADH dehydrogenase subunit 2 |
| KR732847 | *Odontophorus leucolaemus* | NADH dehydrogenase subunit 2 |
| KR732846 | *Odontophorus speciosus* | NADH dehydrogenase subunit 2 |
| KR732845 | *Odontophorus erythrops* | NADH dehydrogenase subunit 2 |
| KR732844 | *Odontophorus guttatus* | NADH dehydrogenase subunit 2 |
| KR732843 | *Odontophorus gujanensis* | NADH dehydrogenase subunit 2 |
| KR732842 | *Odontophorus capueira* | NADH dehydrogenase subunit 2 |
| KR732841 | *Odontophorus stellatus* | NADH dehydrogenase subunit 2 |
| KR732840 | *Dactylortyx thoracicus* | NADH dehydrogenase subunit 2 |
| KR732839 | *Dendrortyx macroura* | NADH dehydrogenase subunit 2 |
| KR732838 | *Callipepla squamata* | NADH dehydrogenase subunit 2 |
| KR732837 | *Callipepla douglasii* | NADH dehydrogenase subunit 2 |
| KC556543 | *Colinus leucopogon* | NADH dehydrogenase subunit 2 |
| KC556066 | *Dendrortyx leucophrys* | NADH dehydrogenase subunit 2 |
| KC556060 | *Cyrtonyx ocellatus* | NADH dehydrogenase subunit 2 |
| KC556524 | *Odontophorus balliviani* | NADH dehydrogenase subunit 2 |
| KC556517 | *Odontophorus columbianus* | NADH dehydrogenase subunit 2 |
| KC556515 | *Odontophorus strophium* | NADH dehydrogenase subunit 2 |
| KC556513 | *Odontophorus melanonotus* | NADH dehydrogenase subunit 2 |
| KC556512 | *Odontophorus hyperythrus* | NADH dehydrogenase subunit 2 |
| KC556507 | *Odontophorus melanotis* | NADH dehydrogenase subunit 2 |
| DQ768288 | *Francolinus nahani* | NADH dehydrogenase subunit 2 |
| KR732851 | *Ptilopachus petrosus* | NADH dehydrogenase subunit 2 |
| EF569482 | *Polyplectron inopinatum* | NADH dehydrogenase subunit 2 |
| EF569481 | *Polyplectron napoleonis* | NADH dehydrogenase subunit 2 |
| EF569480 | *Polyplectron chalcurum* | NADH dehydrogenase subunit 2 |
| EF569479 | *Polyplectron bicalcaratum* | NADH dehydrogenase subunit 2 |
| DQ768268 | *Polyplectron malacense* | NADH dehydrogenase subunit 2 |
| DQ768266 | *Polyplectron germaini* | NADH dehydrogenase subunit 2 |
| KC778823 | *Polyplectron katsumatae* | NADH dehydrogenase subunit 2 |
| KR732830 | *Cyrtonyx montezumae* | 12S ribosomal RNA |
| KR732829 | *Dactylortyx thoracicus* | 12S ribosomal RNA |
| KR732828 | *Oreortyx pictus* | 12S ribosomal RNA |
| KR732827 | *Odontophorus erythrops* | 12S ribosomal RNA |
| KR732826 | *Odontophorus gujanensis* | 12S ribosomal RNA |
| KR732825 | *Odontophorus stellatus* | 12S ribosomal RNA |
| KR732824 | *Odontophorus capueira* | 12S ribosomal RNA |
| KR732823 | *Odontophorus speciosus* | 12S ribosomal RNA |
| KR732822 | *Odontophorus leucolaemus* | 12S ribosomal RNA |
| KR732821 | *Odontophorus balliviani* | 12S ribosomal RNA |
| KR732820 | *Dendrortyx macroura* | 12S ribosomal RNA |
| KR732819 | *Philortyx fasciatus* | 12S ribosomal RNA |
| KR732818 | *Callipepla gambelii* | 12S ribosomal RNA |
| KR732817 | *Callipepla californica* | 12S ribosomal RNA |
| KR732816 | *Callipepla douglasii* | 12S ribosomal RNA |
| KR732815 | *Callipepla squamata* | 12S ribosomal RNA |
| KR732814 | *Colinus cristatus* | 12S ribosomal RNA |
| KR732813 | *Colinus nigrogularis* | 12S ribosomal RNA |
| KR732812 | *Colinus virginianus* | 12S ribosomal RNA |
| KR732832 | *Ptilopachus petrosus* | 12S ribosomal RNA |
| KC749467 | *Polyplectron inopinatum* | 12S ribosomal RNA |
| KC749466 | *Polyplectron germaini* | 12S ribosomal RNA |
| KC749465 | *Polyplectron napoleonis* | 12S ribosomal RNA |
| KC749464 | *Polyplectron chalcurum* | 12S ribosomal RNA |
| KC749463 | *Polyplectron bicalcaratum* | 12S ribosomal RNA |
| KC778974 | *Polyplectron katsumatae* | 12S ribosomal RNA |
| KR732895 | *Colinus cristatus* | eukaryotic elongation factor 2 |
| KR732894 | *Callipepla squamata* | eukaryotic elongation factor 2 |
| KR732893 | *Dendrortyx macroura* | eukaryotic elongation factor 2 |
| KR732892 | *Philortyx fasciatus* | eukaryotic elongation factor 2 |
| KR732891 | *Cyrtonyx montezumae* | eukaryotic elongation factor 2 |
| KR732889 | *Rhynchortyx cinctus* | eukaryotic elongation factor 2 |
| KR732888 | *Oreortyx pictus* | eukaryotic elongation factor 2 |
| KR732887 | *Colinus virginianus* | eukaryotic elongation factor 2 |
| KR732886 | *Callipepla douglasii* | eukaryotic elongation factor 2 |
| KR732885 | *Callipepla californica* | eukaryotic elongation factor 2 |
| KR732884 | *Callipepla gambelii* | eukaryotic elongation factor 2 |
| KR732883 | *Odontophorus erythrops* | eukaryotic elongation factor 2 |
| KR732882 | *Odontophorus guttatus* | eukaryotic elongation factor 2 |
| KR732881 | *Odontophorus speciosus* | eukaryotic elongation factor 2 |
| KR732880 | *Odontophorus leucolaemus* | eukaryotic elongation factor 2 |
| KR732879 | *Odontophorus balliviani* | eukaryotic elongation factor 2 |
| KR732878 | *Odontophorus stellatus* | eukaryotic elongation factor 2 |
| KR732877 | *Odontophorus capueira* | eukaryotic elongation factor 2 |
| KR732890 | *Ptilopachus petrosus* | eukaryotic elongation factor 2 |
| KC749707 | *Polyplectron inopinatum* | eukaryotic elongation factor 2 |
| KC749706 | *Polyplectron germaini* | eukaryotic elongation factor 2 |
| KC749705 | *Polyplectron chalcurum* | eukaryotic elongation factor 2 |
| KC749704 | *Polyplectron bicalcaratum* | eukaryotic elongation factor 2 |
| KC778867 | *Polyplectron katsumatae* | eukaryotic elongation factor 2 |
| KR732875 | *Philortyx fasciatus* | NADH dehydrogenase subunit 5 |
| KR732874 | *Callipepla douglasii* | NADH dehydrogenase subunit 5 |
| KR732872 | *Rhynchortyx cinctus* | NADH dehydrogenase subunit 5 |
| KR732871 | *Dactylortyx thoracicus* | NADH dehydrogenase subunit 5 |
| KR732870 | *Odontophorus leucolaemus* | NADH dehydrogenase subunit 5 |
| KR732869 | *Odontophorus balliviani* | NADH dehydrogenase subunit 5 |
| KR732868 | *Odontophorus guttatus* | NADH dehydrogenase subunit 5 |
| KR732867 | *Odontophorus erythrops* | NADH dehydrogenase subunit 5 |
| KR732866 | *Odontophorus capueira* | NADH dehydrogenase subunit 5 |
| KR732865 | *Odontophorus stellatus* | NADH dehydrogenase subunit 5 |
| KR732864 | *Odontophorus gujanensis* | NADH dehydrogenase subunit 5 |
| KR732863 | *Dendrortyx macroura* | NADH dehydrogenase subunit 5 |
| KR732862 | *Callipepla californica* | NADH dehydrogenase subunit 5 |
| KR732861 | *Callipepla gambelii* | NADH dehydrogenase subunit 5 |
| KR732860 | *Callipepla squamata* | NADH dehydrogenase subunit 5 |
| KR732859 | *Colinus virginianus* | NADH dehydrogenase subunit 5 |
| KR732858 | *Colinus nigrogularis* | NADH dehydrogenase subunit 5 |
| KR732873 | *Ptilopachus petrosus* | NADH dehydrogenase subunit 5 |

Since we are interested in retrieving sequences from GenBank using the accession numbers dataset generated above, we will use the ‘sq.retrieve.indirect()’ function in ‘phruta’. Please note that there are two versions of ‘sq.retrieve.*’ in ‘phruta’. The one that we are using in this tutorial, ‘sq.retrieve.indirect()’, retrieves sequences “indirectly” because it requires a table of accession numbers to be passed as an argument (see the ‘acc.table.retrieve()’ function above). I present the information in this tutorial using ‘sq.retrieve.indirect()’ instead of ‘sq.retrieve.direct()’ because ‘sq.retrieve.indirect()’ is more flexible and robust. Specifically, ‘sq.retrieve.indirect()’ allows users to correct issues prior to downloading/retrieving the sequences. For instance, you can add new sequences, species, populations to the resulting ‘data.frame’ from ‘acc.table.retrieve()’. You could even manually assemble your own dataset of accession numbers to be retrieved using ‘sq.retrieve.indirect()’. Instead, ‘sq.retrieve.direct()’ does its best to directly retrieve GenBank sequences for a target set of taxa and set of gene regions. In short, you should be able to catch errors using ‘sq.retrieve.indirect()’ but mistakes will be harder to spot and fix if you’re using ‘sq.retrieve.direct()’.

We still need to retrieve all the sequences from the accessions table generated using ‘acc.table’. Note that since we have specified ‘download.sqs = FALSE’ in ‘sq.retrieve.indirect()’, the sequences retrieved from GenBank are returned in a list and saved as ‘sqs.downloaded’ in the ‘R’ global workspace. If we were to download the sequences to our working directory using the ‘download.sqs = TRUE’ argument, ‘phruta’ would have written all the resulting ‘fasta’ files into a newly created folder named ‘0.Sequences’ located in our working directory.

~~~
‘“{r}
sqs.downloaded <- sq.retrieve.indirect(acc.table = acc.table, download.sqs = FALSE)
”’
~~~

Now, let’s make sure that we are only including sequences that are reliable and from species that we are actually interested in analyzing. We are going to use the ‘sq.curate()’ function for this. We will provide a list of taxonomic names to filter out incorrect sequences (‘filterTaxonomicCriteria’ argument). For instance, we could simply provide a vector of the genera that we are interested in analyzing. This vector must have a length of ‘1’, with all the target genera being separated with ‘|’ (e.g. ‘“*Callipepla*|*Colinus*|*Dendrortyx*”’ if we were interested in only those three genera). For now, we will assume that all of the species we downloaded are relevant to the analyses (i.e. ‘filterTaxonomicCriteria = [AZ]’). Finally, since we are not downloading anything to our working directory, we need to pass our downloaded sequences (‘sqs.downloaded’ object generated above using the ‘sq.retrieve.indirect()’ function) to the ‘sqs.object’ argument in ‘sq.curate()’.

~~~
‘“{r}
sqs.curated <- sq.curate(filterTaxonomicCriteria = ‘[AZ]’, kingdom = ‘animals’, sqs.object =
sqs.downloaded, removeOutliers = FALSE)
”’
~~~

Running the ‘sq.curate()’ function will create an object of class ‘list’ (i.e. ‘sqs.curated’) that includes (1) the curated sequences with original names, (2) the curated sequences with species-level names (‘renamed_*’ prefix), (3) the accession numbers table (‘AccessionTable’; Table 2), and (4) a summary of taxonomic information for all the species sampled in the files (Table 3). From here, we will align the sequences that we just curated using ‘sq.aln()’ under default parameters. The ‘sq.aln()’ function uses the alignment routines implemented in ‘DECIPHER’ (Wright 2016, 2020) We are again passing the output from ‘sq.curate()’, the object ‘sqs.curated’, using the ‘sqs.object’ argument in ‘sq.aln()’.

~~~
‘“{r}
sqs.aln <- sq.aln(sqs.object = sqs.curated)
”’
~~~

**Table 3.**
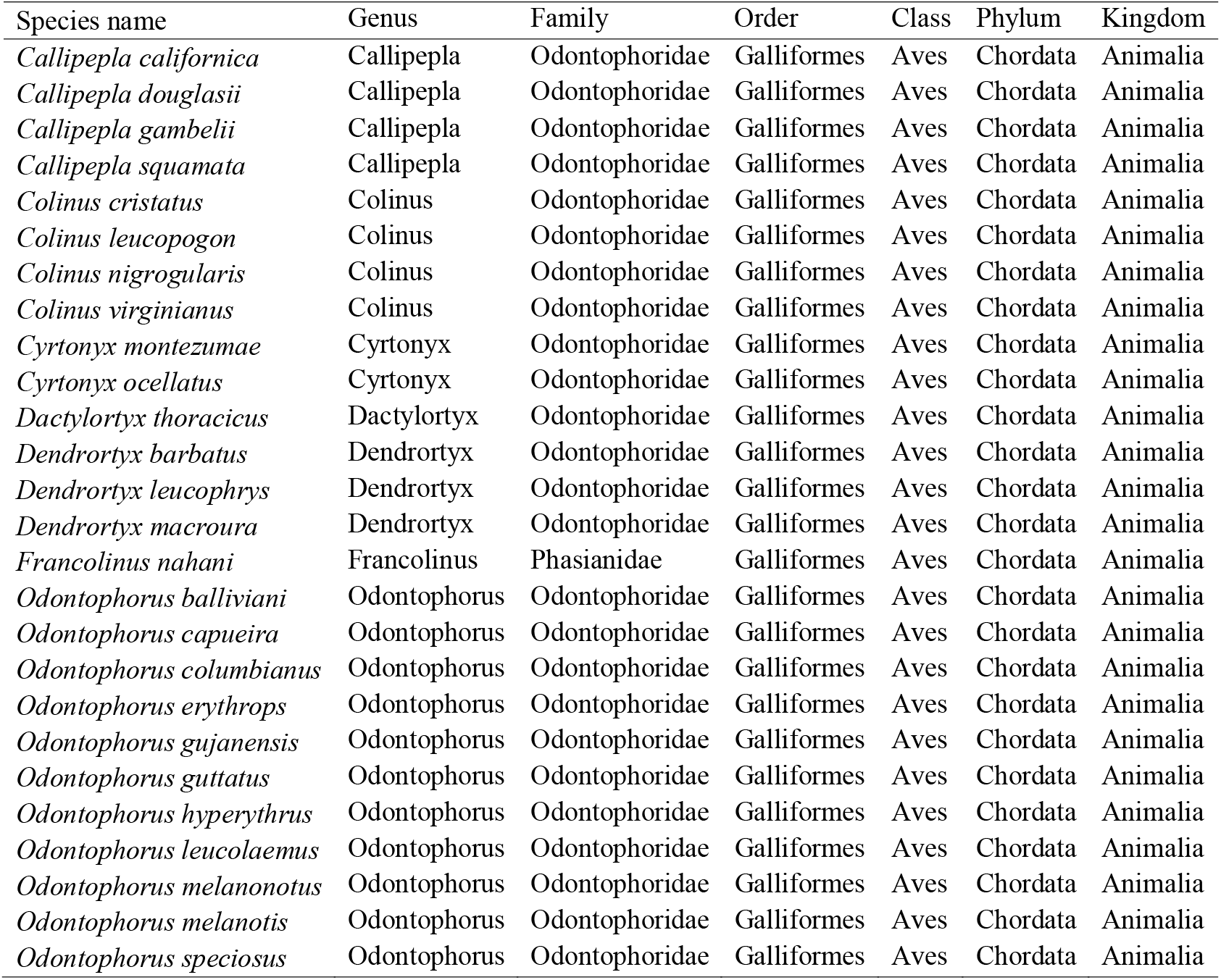
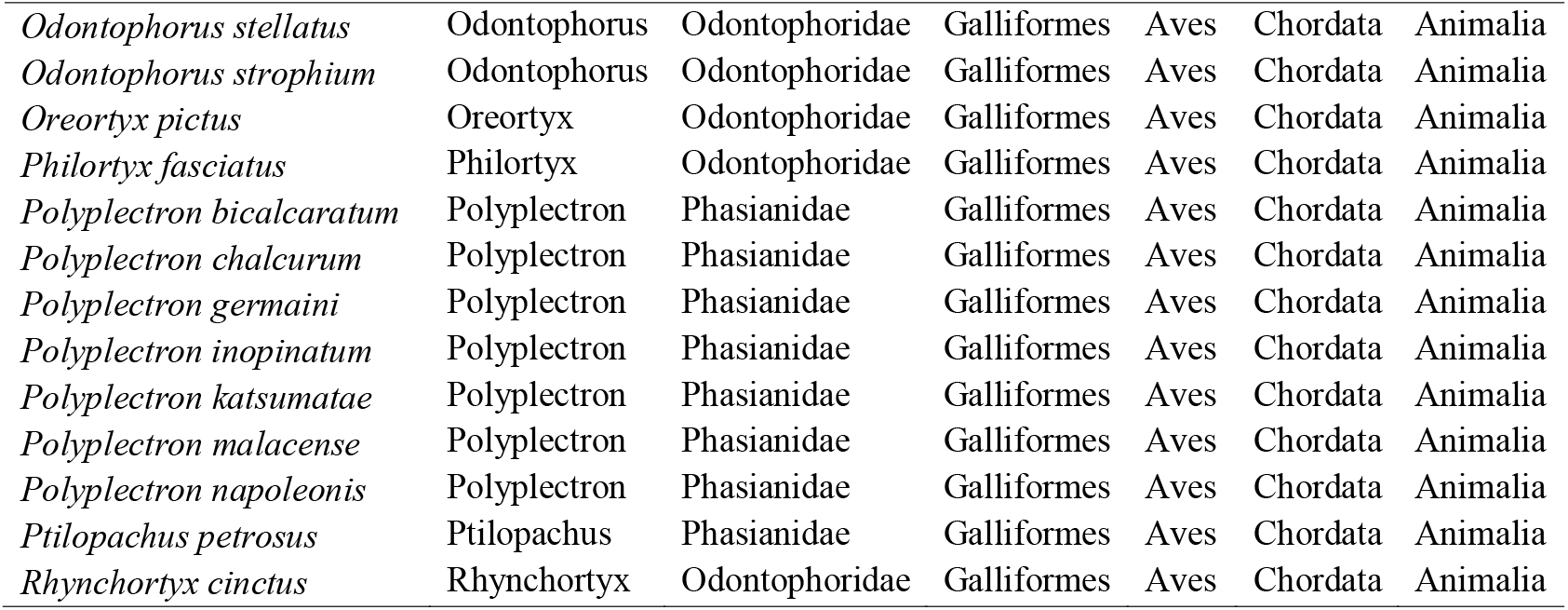
Summary of taxonomic information for the species sampled for Odontophoridae and *Polyplectron* using ‘phruta’. Taxonomic data follows that from the gbif taxonomic backbone for the target species retrieved based on Table 2.

The resulting multiple sequence alignments will be saved to the ‘sqs.aln’ object in our workspace, a list of alignments. For each of the gene regions, we will have access to the original alignment (‘Aln.Original’), the masked one (‘Aln.Masked’), and information on the masking process. The masking process is conducted using the ‘DECIPHER::RemoveGaps()’ function by removing positions where gaps are common to all sequences. The raw and masked alignments are presented in Figures 1 and 2, respectively.

**Figure 1.**
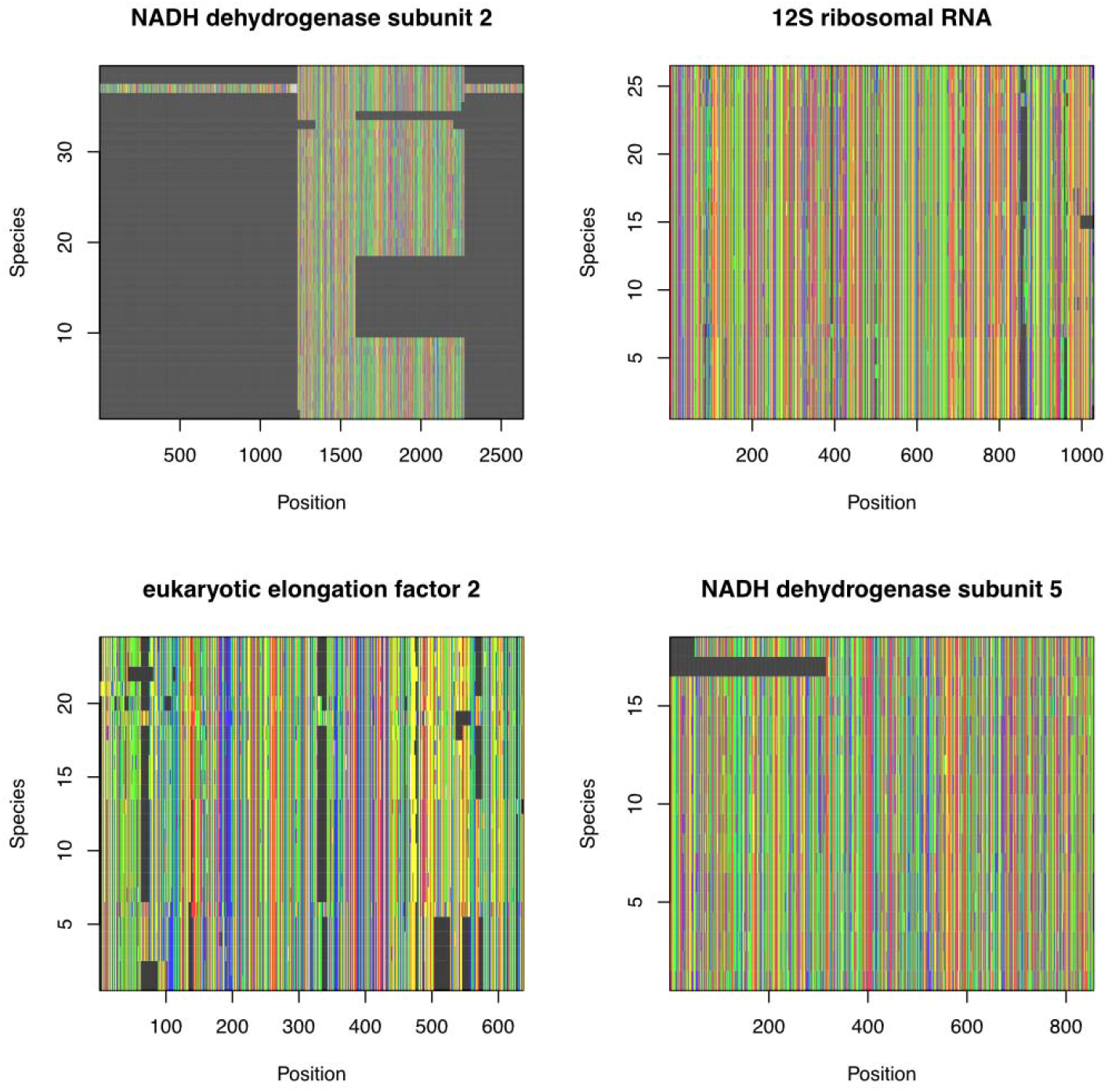
Sequence alignments for the Odontophoridae and *Polyplectron*, the selected outgroup, generated using ‘phruta’. Highly ambiguous sites have not been masked in this figure. Sequence alignments are shown for four genes that were identified as being represented in >20% of the species sampled in GenBank for the target taxa. In each panel, each column represents a site or position in the alignment. Rows reflect species in the relevant dataset. Gray cells represent gaps in the species/position combination.

**Figure 2.**
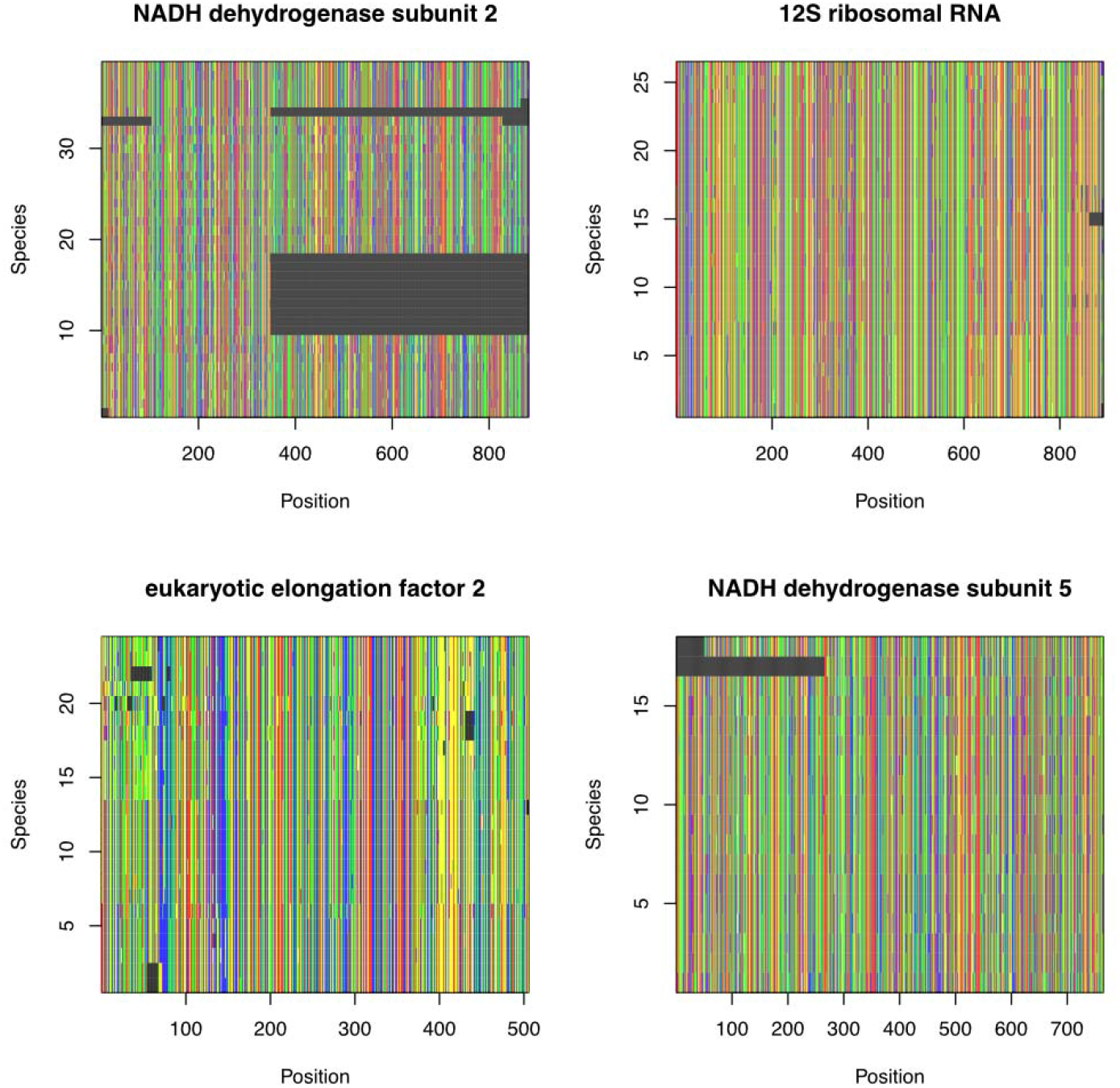
Sequence alignments for the Odontophoridae and *Polyplectron*, the selected outgroup, generated using ‘phruta’. This figure is equivalent to Figure 1 except that highly ambiguous sites have been masked in this figure. Sequence alignments are shown for four genes that were identified as being represented in >20% of the species sampled in GenBank for the target taxa. In each panel, each column represents a site or position in the alignment. Rows reflect species in the relevant dataset. Gray cells represent gaps in the species/position combination.

We have so far generated a molecular dataset for the target clade. Let’s now review how to conduct basic phylogenetic inference in ‘phruta’. Phylogenetic inference in ‘phruta’ is conducted using the ‘tree.raxml()’ function, a wrapper for the ‘ips’ ‘R’ package. To use this function, we will have to export our sequence alignments locally. We will follow the same folder structure as if we were exporting everything locally from ‘phruta’ in each step. Specifically, our sequence alignments must be located in a folder named ‘2.Alignments’ within our working directory. For this tutorial, we will only export the alignments that were masked. We would not need to generate this folder manually if we were already exporting the results from previous functions in ‘phruta’.

~~~
‘‘‘{r}
outgroup <- sqs.curated$Taxonomy[sqs.curated$Taxonomy$genus == ‘Polyplectron’,]
tree.raxml(folder = ‘2.Alignments’,
FilePatterns = ‘Masked_’,
raxml_exec = ‘raxmlHPC’,
Bootstrap = 100,
outgroup = paste(outgroup$species_names, collapse = “,”)
)
’’’
~~~

We are now ready to run RAxML (v.8.2 for this tutorial). Note that in ‘tree.raxml()’, we will need to indicate where the aligned sequences are located (‘folder’ argument), the patterns of the files in the same folder (‘FilePatterns’ argument; “‘Masked_’” in our case), and the total of bootstrap replicates. The ‘outgroup’ argument is optional but since we are interested in calibrating our tree afterwards, we will define it using all the species in *Polyplectron*.

~~~
‘“{r}
outgroup <- sqs.curated$Taxonomy[sqs.curated$Taxonomy$genus == ‘Polyplectron’,]
tree.raxml(folder = ‘2.Alignments’,
       FilePatterns = ‘Masked_’,
       raxml_exec = ‘raxmlHPC’,
       Bootstrap = 100,
       outgroup = paste(outgroup$species_names, collapse = “,”)
       )
”’
~~~

The resulting phylogenetic trees are saved in the ‘3.Phylogeny’ folder, created also in our working directory. For many, the bipartitions tree generated in these runs, ‘RAxML_bipartitions.phruta’, are among the most relevant files (Fig. 3). The ‘3.Phylogeny’ folder further includes the additional ‘RAxML’-related input and output files. Note that users can also run partitioned analyses in ‘RAxML’ within ‘phruta’. This approach is available by setting the ‘partitioned’ argument in ‘tree.raxml()’ to ‘TRUE’. For now, partitioned analyses are based on the gene-based regions that are being analyzed. The same model is used to analyze each partition. More details on partitioned analyses can be customized by passing arguments in ‘ips::raxml()’.

~~~
‘“{r}
tree.raxml(folder = “2.Alignments”, FilePatterns = “Masked_”,
       raxml_exec = “raxmlHPC”, Bootstrap = 100,
       outgroup = paste(outgroup$species_names, collapse = “,”),
       partitioned = TRUE
       )
”’
~~~

**Figure 3.**
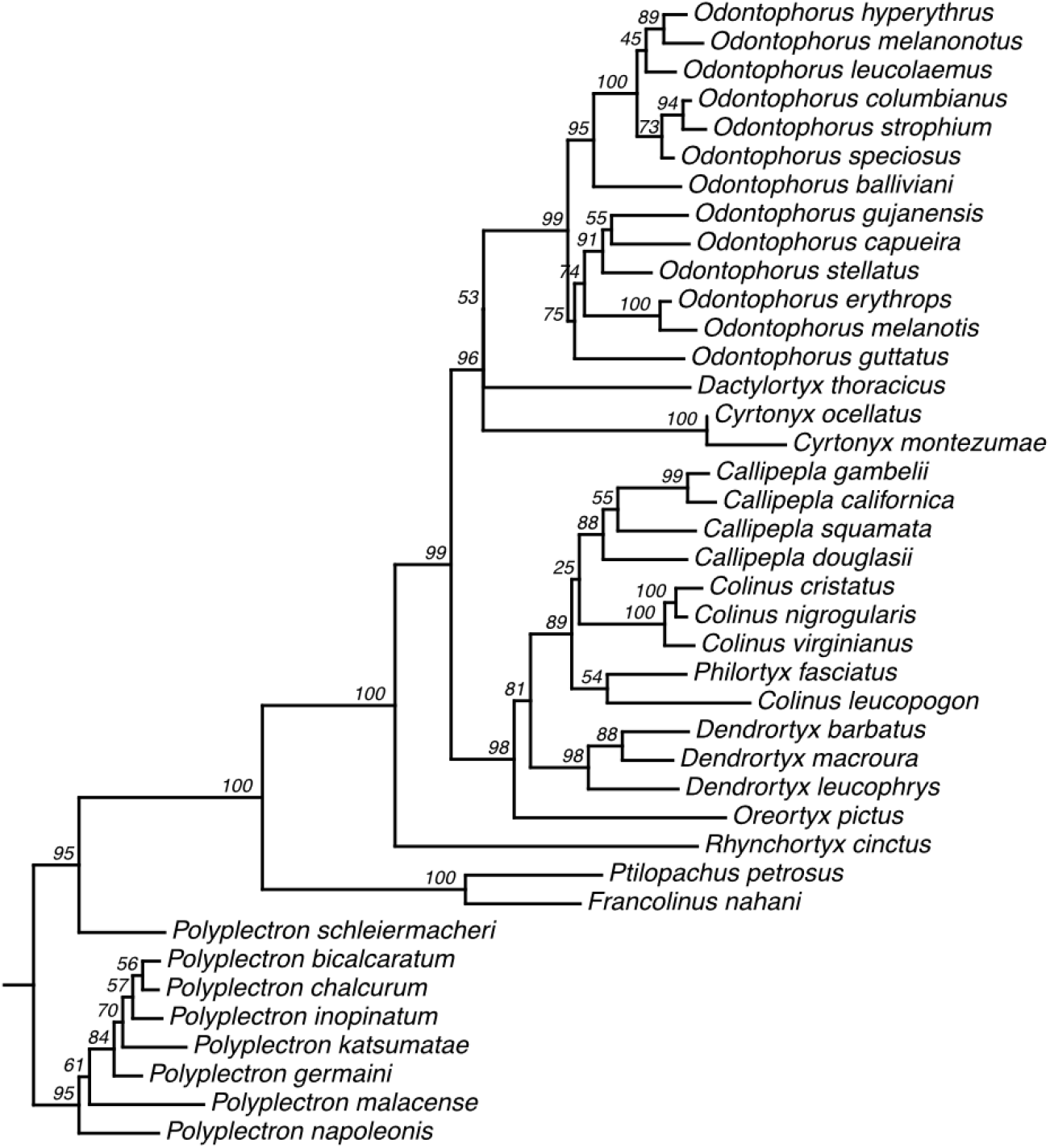
Phylogenetic relationships for the Odontophoridae and *Polyplectron*, the selected outgroup, generated using ‘phruta’ and ‘RAxML’. The bipartitions tree is shown in figure, with support values based on a total of 1,000 bootstrap replicates. This tree was constructed using a concatenated (but unpartitioned) alignment of all four gene alignments presented in Figure 2.

Finally, we note that users will sometimes need to generate constraint trees to seed their searches or limit the tree sampling space. These constraints can be generated using other software such as TACT (Chang et al. 2019).

## 4 TREE DATING IN ‘phruta’

‘phruta’ includes a basic wrapper for functions included in the ‘geiger’ ‘R’ package that allows for the time-calibration of target phylogenies. Based on the tutorial presented above, we can perform basic tree dating of our phylogeny using secondary calibrations extracted from Scholl and Wiens (2016) phylogeny. I am only using this study because it has a large phylogeny with detailed taxonomic information available for each terminal. I expect to include additional trees in the near future. Users can also choose to use their own reference phylogeny if available. Note that the ‘tree.dating()’, the function in ‘phruta’ that is able to conduct phylogenetic dating based on node correspondence between phylogenies, requires the user to specify where the ‘1.Taxonomy.csv’ file is located. This file, ‘1.Taxonomy.csv’, is created automatically when sequences are curated using ‘sq.curate()’ and results are exported into your local repository. However, since we have been keeping our results in the ‘R’ global environment, we will have to export ‘1.Taxonomy.csv’ manually before we can move forward and time-calibrate the tree.

~~~
‘“{r}
dir.create(“1.CuratedSequences”)
write.csv(sqs.curated$Taxonomy, ‘1.CuratedSequences/1.Taxonomy.csv’)
”’
~~~

Tree dating is performed using the ‘tree.dating()’ function in ‘phruta’. For this function, we have to provide the name of the folder containing the ‘1.Taxonomy.csv’ file created in ‘sq.curate()’. We also have to indicate the name of the folder containing the ‘RAxML_bipartitions.phruta’ file. We will scale our phylogeny using ‘treePL’.

~~~
‘“{r}
tree.dating(taxonomyFolder = “1.CuratedSequences”,
        phylogenyFolder = “3.Phylogeny”,
        scale = ‘treePL’)
”’
~~~

Running this line will result in a new folder ‘4.Timetree’, which includes the different time-calibrated phylogenies obtained (if any) and associated secondary calibrations used in the analyses. The resulting time-calibrated tree for the analyses presented in this article is presented in Figure 4.

**Figure 4.**
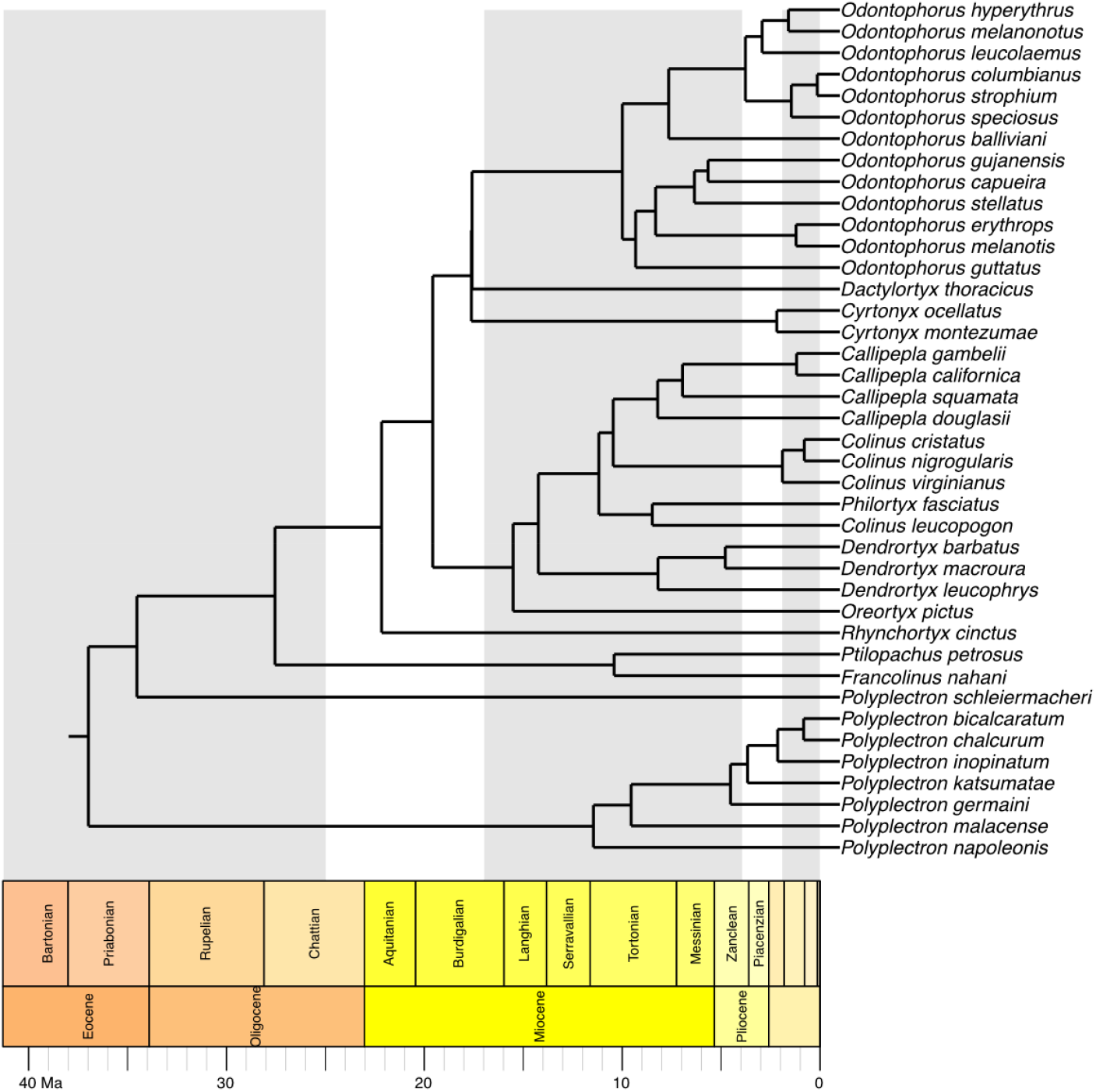
Time-calibrated phylogeny for the Odontophoridae and *Polyplectron*, the selected outgroup, generated using ‘phruta’ and ‘RAxML’ and secondary calibrations. The backbone topology is presented in Figure 5. The same figure shows support values.

## 5 ADVANCED METHODS WITH ‘phruta’

### 5.1 Curating taxonomic names

You can use ‘taxonomy.retrieve()’, a function implemented inside ‘sq.curate()’ in ‘phruta’ to curate species names. For instance, the block of code below will curate taxonomic names using the gbif backbone taxonomy (GBIF Secretariat, 2022). Note that the ‘kingdom’ argument in ‘taxonomy.retrieve()’ can be set to ‘NULL’, meaning that there will be no indication of the kingdom of the taxa when performing taxonomic searches.

~~~
‘“{r}
phruta:::taxonomy.retrieve(species_names=c(“Felis_catus”, “PREDICTED:_Vulpes”,
            “Phoca_largha”, “PREDICTED:_Phoca”,
            “PREDICTED:_Manis”, “Felis_silvestris”, “Felis_nigripes”),
            database = ‘gbif)
”’
~~~

However, ‘gbif is efficient for retrieving accurate taxonomy when we provide details on the ‘kingdom’. In the same example, and given that all the species we are interested in are animals, we could just use the following block of code to curate taxonomic names.

~~~
‘“{r}
phruta:::taxonomy.retrieve(species_names = c(“Felis_catus”, “PREDICTED:_Vulpes”,
            “Phoca_largha”, “PREDICTED:_Phoca”,
            “PREDICTED:_Manis”, “Felis_silvestris”, “Felis_nigripes”),
            database = ‘gbif’, kingdom = ‘animals’)
”’
~~~

Depending on your sampling, you could also do the same for plants by using ‘plants’ in the ‘kingdom’ argument instead of ‘animals’. Now, what if we were interested in following other databases to retrieve taxonomic information for the species in our database? The latest version of ‘phruta’ allows users to select the desired database. The databases follow the ‘taxize::classification()’ function (Chamberlain and Szöcs, 2013).

~~~
‘“{r}
phruta:::taxonomy.retrieve(species_names = c(“Felis_catus”, “PREDICTED:_Vulpes”,
            “Phoca_largha”, “PREDICTED:_Phoca”,
            “PREDICTED:_Manis”, “Felis_silvestris”, “Felis_nigripes”),
            database = ‘itis’)
”’
~~~

### 5.2 Running PartitionFinder in ‘phruta’

With the current version of ‘phruta’, users are able to run ‘PartitionFinder’ v1 (Lanfear et al. 2012) from ‘R’. For this, users should provide the name of the folder where the alignments are stored, a particular pattern in the file names (‘Masked_’ in our case), and which models will be run in ‘PartitionFinder’. This function will download ‘PartitionFinder’, generate the input files, and run the software, all within R. The output files will be in a new folder within the working directory.

~~~
‘“{r}
sq.partitionfinderv1(folderAlignments = “2.Alignments”,
           FilePatterns = “Masked_”,
           models = “all”
           )
”’
~~~

Unfortunately, the output files are not integrated with the current ‘phruta’ pipeline. This will be part of a new release. However, users can still perform gene-based partitioned analyses within ‘RAxML’ or use ‘PartitionFinder’’s output files to inform their own analyses outside ‘phruta’. An upcoming release of ‘phruta’ will allow users to run more recent versions of PartitionFinder.

### 5.3 Identifying rogue taxa

‘phruta’ can help users run ‘RogueNaRok’ (Aberer et al. 2011) implemented in the ‘Rogue’ R package (Aberer et al. 2013). Users can examine whether rogue taxa should be excluded from the analyses. ‘tree.roguetaxa()’ uses the bootstrap trees generated using the ‘tree.raxml()’ function along with the associated best tree to identify rogue taxa.

~~~
‘“{r}
tree.roguetaxa(folder = “3.Phylogeny”)
”’
~~~

## 6 REPRODUCIBILITY WITH ‘phruta’

One of the central points for developing ‘phruta’ was related to increasing the reproducibility of relatively simple phylogenetic analyses. By either compiling or calling alternative tools that are commonly used to assemble species-level molecular and phylogenetic datasets inside an ‘R’ package, ‘phruta’ allows users to generate a clear, structured, and reproducible workflow. In fact, ‘phruta’ is conceived as a package that allows users to choose between at least two alternative options to share their workflow. First, users can simply provide access to their workflow in an ‘R’ script. For instance, this file can be stored in GitHub along with all the intermediate files that are created at each given step. Alternatively, given that the information in databases is constantly changing, users can share their ‘R’ script and associated workspace to assure that the versions of the retrieved files correspond to specific versions of the databases.

As an example, an early version of this manuscript, written in RMarkdown (Baumer and Udwin, 2015), is available from https://github.com/cromanpa94/phruta_ms. This ‘RMarkdown’ will allow users to replicate the analyses presented in the current paper. With the workspace, saved from the RMarkdown as

~~~
‘“{r}
save.image(file = “phruta_ms.RData”)
”’
~~~

, users can choose to use the stored objects instead of performing searches on GenBank and additional taxonomic databases. These two files confer increased reproducibility to the analyses presented in this article.

## 7 PERFORMANCE

Assessing the performance of ‘phruta’ is intrinsically challenging. However, it is expected for functions that scrape and curate information stored in GenBank to show a slow performance on search terms (e.g. clades) with extensive genetic sampling. Below, I focus on comparing the distribution of durations for assembling molecular datasets in the analyses presented in Section 3 in the current document. These estimates encompass all the steps outlined above ranging between using the functions ‘gene.sampling.retrieve()’ and ‘sq.curate’. In particular, performance analyses focused on defining the number of simultaneous searches conducted in ‘phruta’ and the number of examined hits analyzing each time from GenBank. Time estimates were calculated using the ‘microbenchmark’ ‘R’ package (Mersmann 2021).

Figure 5 shows a summary of ‘phruta’ performance under different search parameters for ‘gene.sampling.retrieve()’ and ‘sq.curate’. Overall, assembling and curating the analyses presented in Section 3 took consistently less than 6 minutes to complete regardless of the parameters used to define searches in ‘phruta’. As expected, the more simultaneous searches conducted, the larger the decrease in computational time. Similarly, the greater the number of GenBank hits examined per search, the lower the computational time. These differences related to the number of hits examined are inversely proportional to the number of cores used during the search process. Finally, note that, except when using 8 simultaneous cores, searches are all completed successfully. When 8 cores are used, only searches for small batches of hits (n=50 or 100) are successful. We used these patterns to define the optimal (i.e. fastests and successful) search parameters shown in this paper and used for default in ‘phruta’ and ‘salphycon’.

**Figure 5.**
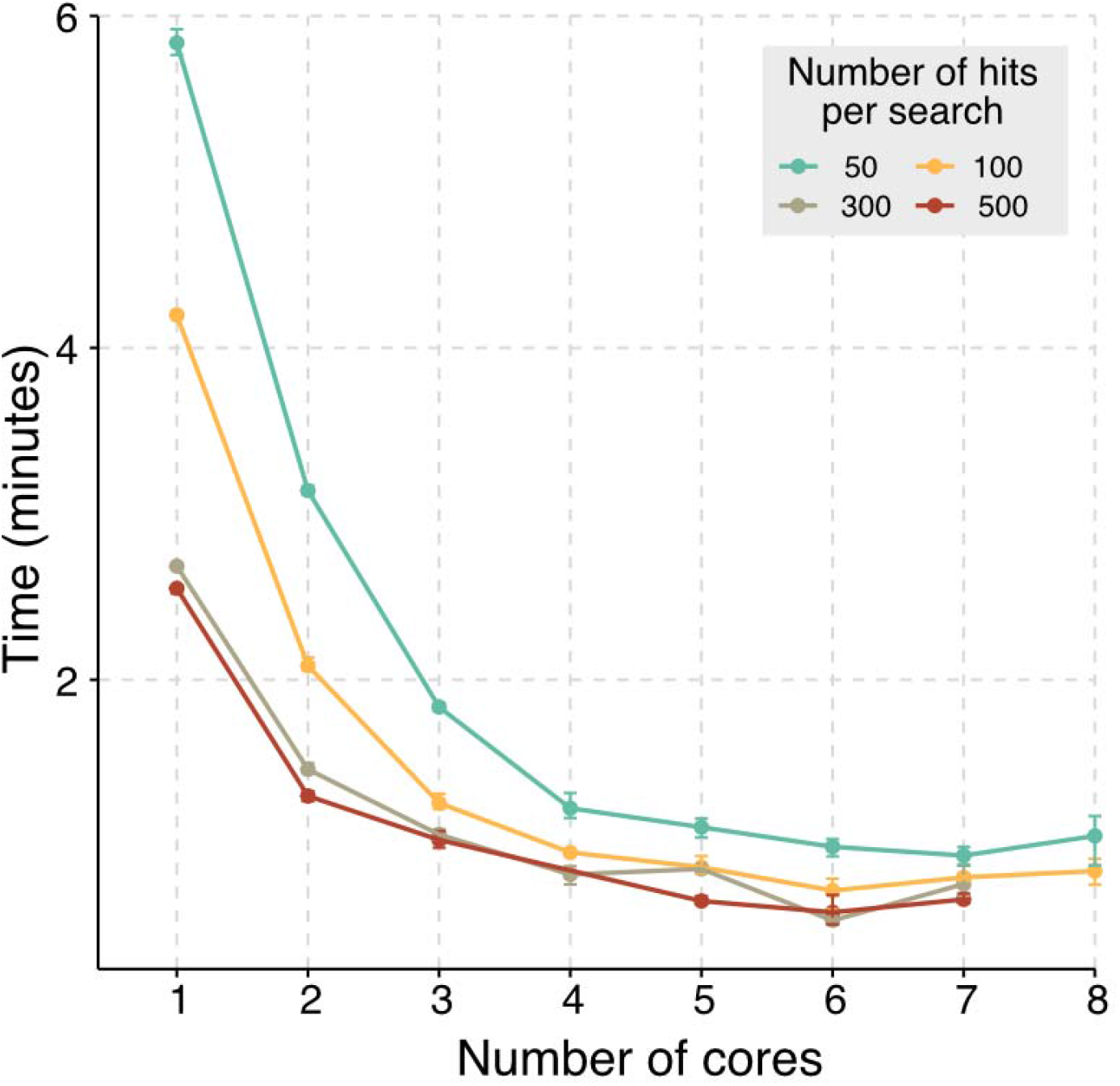
Results for performance analyses showing the duration of searches conducted based on the combination of number of hits per search (i.e. number of hits analyzed each time the search is conducted) and the number of simultaneous searches conducted from ‘phruta’. Note that not all combinations produced results. The search parameters used in this article were optimized based on this analysis.

## 8 THE ‘salphycon’ WEB APP

To support the use of ‘phruta, I developed ‘salphycon’, a web app that extends the basic functionality of ‘phruta’ into a graphical user interface (Figure 6). The app written in Shiny, ‘salphycon’, allows users to run the fundamental functions in ‘phruta’ without the need of writing code. The app also extends ‘phruta’ to Spanish-speaking users by including translations in the same language using the ‘shi18ny’ ‘R’ package (Marin Diaz, 2022).

**Figure 6.**
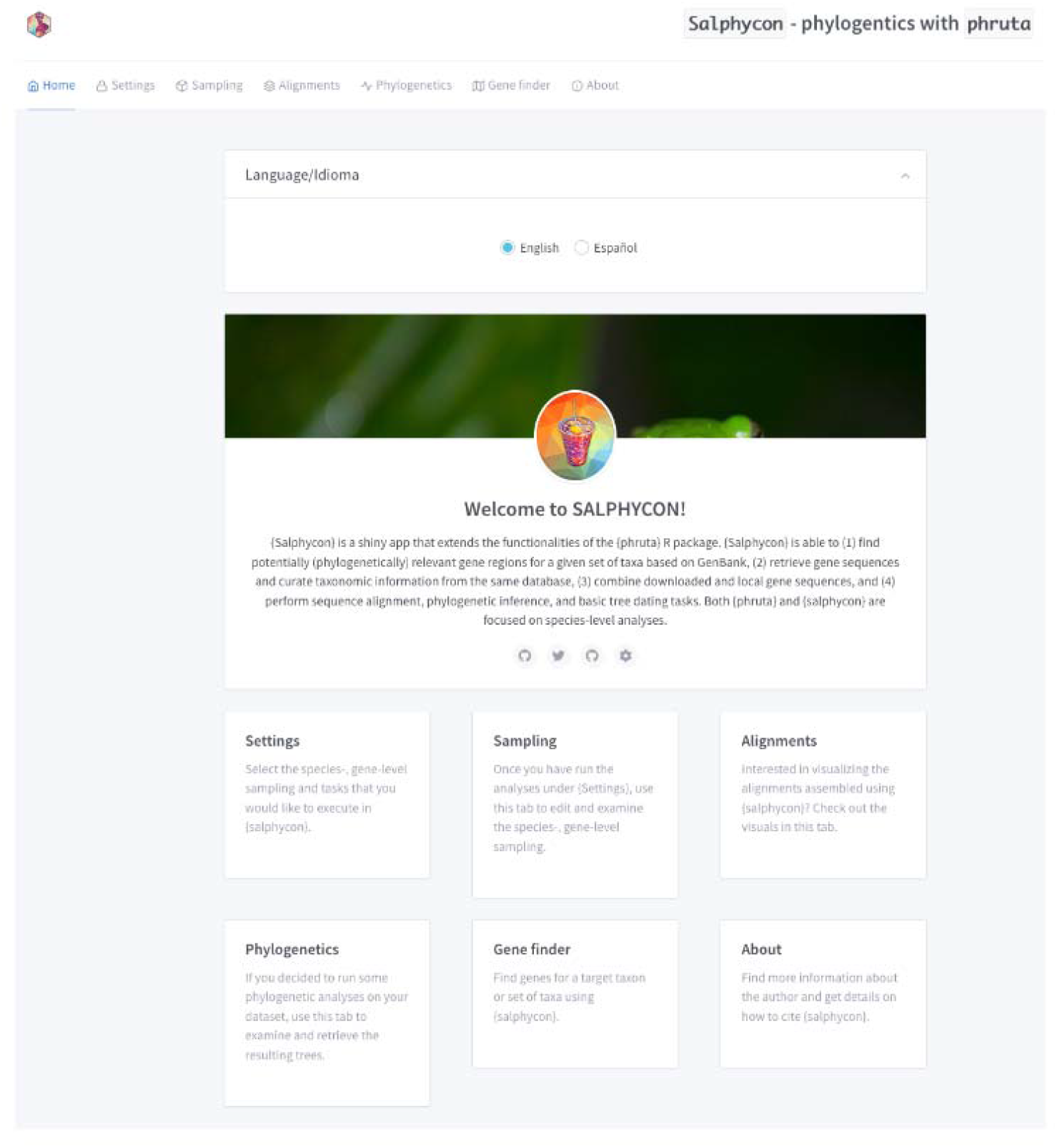
Landing page for ‘salphycon’, the web application including a set of the basic functions implemented in ‘phruta’. In this app, users are able to retrieve sequences from GenBank, explore sequence alignments, and conduct basic phylogenetic inference with RAxML. Users can also choose to define the language of the app.

## 9 CONCLUSIONS

Both ‘phruta’ and ‘salphycon’ were developed to increase access and reproducibility in phylogenetic analyses. These two tools, released under open access code, following best coding practices, and with transparent implementation, extend the functionality of existing libraries in R and additional software that is commonly used in the phylogenetic workflow. Importantly, ‘phruta’ and ‘salphycon’ enable the exploration of existing gene regions and sequences deposited in GenBank to be used in phylogenetic analyses. We note, however, that both of these tools are limited in different ways. Current limitations of ‘phruta’ include (1) the assemblage of phylogenies with sequences at the species level, (2) the inference of of single-gene or multi-locus phylogenies instead of phylogenomics, (3) the implementation of PartitionFinder v1 instead of the latest release (Lanfear et al. 2017), (4) intrinsic limitations associated with the quality of data hosted in GenBank, (5) the lack of flexibility for choosing among more commonly used alignment and phylogenetic packages, among others. Future releases intend to extend the toolbox in ‘phruta’ to overcome these and other limitations and to explore the implementation of utilities for phylogenetic comparative methods within ‘salphycon’.

## AUTHORS CONTRIBUTIONS’

C.R.P. wrote the R package, shiny app, vignettes, documentation, developed the associated testing framework, and wrote the paper.

## ACKNOWLEDGEMENTS

The author thanks Heidi E. Steiner for proofreading the vignettes and documentation in ‘phruta’ in addition to early versions of this manuscript. Anna Krystalli, Rayna Harris, Frederick Boehm, and Maëlle Salmon provided excellent comments during the peer review process of ‘phruta’ in ROpenSci. Wonkyiun Yim provided early support in ‘salphycon’. Finally, the author thanks the author and maintainers of all the packages used in ‘phruta’ and ‘salphycon’ for their constant support to the field.

## DATA AVAILABILITY STATEMENT

The ‘phruta’ R package code is open-source and available on github at https://github.com/ropensci/phruta. The associated shiny app, ‘salphycon’, is also available from GitHub at https://github.com/cromanpa94/salphycon. An early version of this manuscript, which is fully reproducible, is available at https://github.com/cromanpa94/phruta_ms.

